# A *w*Mel *Wolbachia* variant in *Aedes aegypti* from field-collected *Drosophila melanogaster* with increased phenotypic stability under heat stress

**DOI:** 10.1101/2022.01.02.474744

**Authors:** Xinyue Gu, Perran A. Ross, Julio Rodriguez-Andres, Katie L. Robinson, Qiong Yang, Meng-Jia Lau, Ary A. Hoffmann

**Affiliations:** Pest and Environmental Adaptation Research Group, Bio21 Institute and the School of BioSciences, The University of Melbourne, Parkville, Victoria, Australia; Peter Doherty Institute for Infection and Immunity and Faculty of Veterinary and Agricultural Sciences, The University of Melbourne, VIC 3000 Melbourne, Australia

**Keywords:** *Aedes aegypti*, *Wolbachia*, microinjection, heat tolerance, *w*Mel

## Abstract

Mosquito-borne diseases such as dengue, Zika and chikungunya remain a major cause of morbidity and mortality across tropical regions. Population replacement strategies involving the *w*Mel strain of *Wolbachia* are being used widely to control mosquito-borne diseases transmitted by *Aedes aegypti*. However, these strategies may be influenced by environmental temperature because *w*Mel is vulnerable to heat stress. *w*Mel infections in their native host *Drosophila melanogaster* are genetically diverse, but few transinfections of *w*Mel variants have been generated in *Ae. aegypti* mosquitoes. Here we successfully transferred a *w*Mel variant (termed *w*MelM) originating from a field-collected *D. melanogaster* population from Victoria, Australia into *Ae. aegypti*. The new *w*MelM variant (clade I) is genetically distinct from the original *w*Mel transinfection (clade III) generated over ten years ago, and there are no genomic differences between *w*MelM in its original and transinfected host. We compared *w*MelM with *w*Mel in its effects on host fitness, temperature tolerance, *Wolbachia* density, vector competence, cytoplasmic incompatibility and maternal transmission under heat stress in a controlled background. *w*MelM showed a higher heat tolerance than *w*Mel, with stronger cytoplasmic incompatibility and maternal transmission when eggs were exposed to heat stress, likely due to higher overall densities within the mosquito. Both *w*Mel variants had minimal host fitness costs, complete cytoplasmic incompatibility and maternal transmission, and dengue virus blocking under standard laboratory conditions. Our results highlight phenotypic differences between closely related *Wolbachia* variants. *w*MelM shows potential as an alternative strain to *w*Mel in dengue control programs in areas with strong seasonal temperature fluctuations.

## Introduction

*Aedes aegypti* mosquitoes transmit some of the most important arboviral diseases such as dengue, which remain a major cause of morbidity and mortality across tropical regions (Kyle and Harris, 2008; Guzman *et al*., 2010). A promising approach to reduce mosquito-borne disease involves the release of *Ae. aegypti* infected with the bacterium *Wolbachia* into wild populations (Hoffmann *et al*., 2011; Garcia *et al*., 2016; Indriani *et al*., 2020; Ahmad *et al*., 2021; Wang *et al*., 2021). *Wolbachia* are common intracellular bacteria that are transmitted maternally and have a range of effects on their insect hosts (Hoffmann and Turelli, 1997). *Wolbachia* often affect the reproduction of their hosts, particularly by causing cytoplasmic incompatibility (Hoffmann and Turelli, 1997), a phenomenon that results in sterility when a *Wolbachia*-infected male mates with an uninfected female or a female carrying a different infection (Caspari and Watson, 1959; Hoffmann and Turelli, 1997). Cytoplasmic incompatibility can be applied directly to suppress field mosquito populations through male-only releases (Zheng *et al*., 2019; Crawford *et al*., 2020; Beebe *et al*., 2021; Ng and Consortium, 2021) or replace populations with mosquitoes carrying *Wolbachia* (Hoffmann *et al*., 2011; Yen and Failloux, 2020). Replacement strategies are undertaken because *Wolbachia* reduce the transmission of dengue and other arboviruses by *Ae. aegypti* (Moreira *et al*., 2009; van den Hurk *et al*., 2012). Effective virus suppression is related to high *Wolbachia* infection frequencies in populations and high densities within individual mosquitoes (Lu *et al*., 2012; Pinto *et al*., 2021).

Through embryo microinjection, several *Wolbachia* infections including *w*Mel, *w*MelPop, *w*MelCS and *w*AlbB have been established in *Ae. aegypti* following interspecific transfers from *Drosophila, Culex* and *Aedes* (Xi *et al*., 2005; McMeniman *et al*., 2009; Walker *et al*., 2011; Fraser *et al*., 2017). Releases to replace existing populations with those carrying *Wolbachia* have been achieved with both the *w*Mel and *w*AlbB strains (Hoffmann *et al*., 2011; Nazni *et al*., 2019; Tantowijoyo *et al*., 2020). *w*Mel originates from *D. melanogaster* (Hoffmann, 1988; Walker *et al*., 2011) and is now widely used for population replacement in *Ae. aegypti,* with releases undertaken in at least 10 countries including Australia, Brazil, Vietnam and Indonesia (Hoffmann *et al*., 2011; Garcia *et al*., 2019; Tantowijoyo *et al*., 2020; Hien *et al*., 2021). Mosquitoes with the *w*AlbB infection have been released in Malaysia, Australia, USA and Singapore for either population replacement or suppression (Nazni *et al*., 2019; Staunton *et al*., 2019; Crawford *et al*., 2020; Beebe *et al*., 2021; Ng and Consortium, 2021). There is now evidence of reduced dengue transmission after *Wolbachia* releases following replacement with both *w*Mel and *w*AlbB (Nazni *et al*., 2019; Ryan *et al*., 2019; Indriani *et al*., 2020; Utarini *et al*., 2021).

The long-term success of *Wolbachia* population replacement strategies will depend on the stability of *Wolbachia* infections in populations and their phenotypic effects (Bull and Turelli, 2013; Hoffmann *et al*., 2015; Ross *et al*., 2019a). One of these factors is the ability to maintain *Wolbachia* effects under fluctuating temperatures. Compared to the range that their hosts can tolerate, *Wolbachia* infections may be more vulnerable to temperature extremes (Corbin *et al*., 2017; Ross *et al*., 2017). However, *Wolbachia* strains in *Ae. aegypti* respond differently to heat stress, with *w*AlbB-infected mosquitoes maintaining stronger cytoplasmic incompatibility, maternal transmission and *Wolbachia* density under heat stress compared to *w*Mel (Ross *et al*., 2017). In addition, the lack of selection response in *w*Mel for increased heat resistance (Ross and Hoffmann, 2018) could limit the suitability of this strain in locations with extreme temperature fluctuations. Cold temperatures and long-term quiescence of eggs can also reduce *Wolbachia* densities, which may lead to reduced virus blockage or cytoplasmic incompatibility following a dry season (Lau *et al*., 2020; Lau *et al*., 2021).

The *w*Mel strain is genetically diverse in natural *D. melanogaster* populations, with six major intraspecific clades (Richardson *et al*., 2012; Early and Clark, 2013; Ilinsky, 2013). Patterns of *w*Mel clades have shifted in recent decades, with the global spread of the *w*Mel variant (clade III) replacing many *w*Mel-CS-like (clade VI) variants (Riegler *et al*., 2005). *w*Mel variants have divergent phenotypic effects on hosts, inducing different levels of protection against viruses (Chrostek *et al*., 2013), expression of cytoplasmic incompatibility (Veneti *et al*., 2003) and fitness costs (Chrostek *et al*., 2013). *Wolbachia* titres also vary across lines and populations which may relate to *w*Mel variants (Chrostek *et al*., 2013; Early and Clark, 2013). The incidence of these variants in *D. melanogaster* is also affected by environmental conditions, with lower *Wolbachia* infection frequencies in cooler temperate regions (Hoffmann, 1988; Kriesner *et al*., 2016). Temperature influences the phenotypic effects of *w*Mel, including levels of virus protection (Chrostek *et al*., 2021) and female fertility (Kriesner *et al*., 2016). However, these different environmental responses may vary among *w*Mel and *w*Mel-like variants as indicated by impacts of variants on maternal transmission under cool temperatures (Hague *et al*., 2022), host temperature preference (Truitt *et al*., 2019; Hague *et al*., 2020a) and survival following heat stress (Gruntenko *et al*., 2017; Burdina *et al*., 2021).

As *w*Mel that has so far been transfected into *Ae. aegypti* is vulnerable to heat stress, and *w*Mel variants in *D. melanogaster* have different thermal responses, we were motivated to generate new *w*Mel variants in *Ae. aegypti* which could show greater environmental stability. Here we report the generation and characterization of an *Ae. aegypti* line (termed *w*MelM) infected with a *w*Mel variant from *D. melanogaster* collected near Melbourne, Australia. We compared *w*MelM to the original *w*Mel strain (generated by Walker *et al*., 2011) in terms of life history, quiescent egg viability, *Wolbachia* density, vector competence and cytoplasmic incompatibility and maternal transmission under cold and heat stress. We found *w*MelM had higher heat tolerance without an obvious fitness cost while maintaining dengue blocking, providing a new strain that may be useful for release in different contexts.

## Methods

### Insect strains and colony maintenance

*Aedes aegypti* mosquitoes were reared in temperature-controlled insectaries at 26 ± 1 °C with a 12 h photoperiod following Ross et al. (2017). Larvae were reared in trays filled with 4L of reverse osmosis (RO) water and provided with fish food (Hikari tropical sinking wafers, Kyorin food, Himeji, Japan) *ad libitum* throughout their development. Four to five hundred adults were maintained in BugDorm-1 (27 L) cages (Megaview Science C., Ltd., Taichung, Taiwan). Four-to-six-day old female mosquitoes were starved for 24 hr, then fed on the forearm of a human volunteer. Blood feeding of female mosquitoes on human volunteers was approved by the University of Melbourne Human Ethics Committee (approval 0723847). All adult subjects provided informed written consent (no children were involved).

Three *Ae. aegypti* populations were used in this study, with an additional population (*w*AlbB) used in DENV vector competence experiments. *w*Mel-infected mosquitoes were collected from Cairns, Australia in 2013 from areas where *w*Mel had established in the wild population (Hoffmann *et al*., 2011) following release of the *w*Mel strain developed by Walker *et al*. (2011). The strain was developed with the shell vial technique which was used to infect the RML-12 cell line with *w*Mel *Wolbachia* from *D. melanogaster* embryos from a laboratory stock (*yw^67C23^*) and maintained by continuous serial passage. *Wolbachia* was then purified from this cell line and injected into *Ae. aegypti* embryos to transfer the *w*Mel infection (Walker *et al*., 2011). In contrast, the *w*MelM-infected mosquitoes in our study were generated by transferring cytoplasm directly from field-collected *D. melanogaster* through microinjection (see *embryonic microinjection* below). *w*Mel.tet mosquitoes were cured of their *w*Mel infection by treating adults with 2 mg/mL tetracycline hydrochloride in a 10% sucrose solution across two consecutive generations. *w*AlbB-infected mosquitoes were generated through microinjection as described previously (Ross *et al*., 2021). Before experiments commenced, all populations were backcrossed to an uninfected population collected from Cairns, Australia for three consecutive generations to control for genetic background.

To generate a line for source material for microinjection for the current study, *D. melanogaster* were collected from the Yarra Valley, Victoria, Australia in April 2019. Isofemale lines were established and their progeny were screened for *Wolbachia* infection (see *Wolbachia* detection and density), with only eggs from *Wolbachia*-infected lines used in experiments. Flies were maintained on cornmeal media 19 ± 1 °C with a 12 h photoperiod (Hoffmann *et al*., 1986).

### Embryonic microinjection

*w*MelM from *D. melanogaster* was introduced to *Ae. aegypti* through embryonic microinjection. Cytoplasm was removed from approximately 1 hr old donor eggs and injected directly into the recipient embryo according to Xi *et al*. (2005). Recipient eggs were collected by placing cups filled with larval rearing water and lined with filter paper (diameter 90 mm) into cages of mosquitoes blood fed >4 d prior. Eggs (<1.5 hr old, light gray in color) were lined up on filter paper and transferred to a cover slip with double-sided tape. Eggs were covered in 100% halocarbon oil and injected using a MINJ-1000 microinjection system (Tritech Research, Los Angeles, CA, USA). Eggs were gently removed from the oil after 10 minutes using a fine paintbrush and placed on a moist piece of filter paper. Eggs were hatched 3 d post-injection by submerging filter papers in containers filled with RO water, a few grains of yeast and half tablet of fish food. Hatching larvae were reared to adulthood and F_0_ females were crossed to uninfected males, blood fed and isolated for oviposition. We screened the females for *w*MelM infection after producing viable offspring using LAMP (Goncalves *et al*., 2019; Jasper *et al*., 2019). This crossing process was repeated for three further generations, with only progeny from *Wolbachia*-positive females contributing to the next generation. We stopped the backcrossing and the detection once the *w*MelM infection reached fixation.

### Whole genome sequencing

We sequenced the whole *Wolbachia* genome of the *w*MelM transinfection generated in this study at F_3_ post-transinfection. We also sequenced populations from three lines at F_7_, which were derived from three single F_1_ females. Genomic DNA was extracted from a pooled sample of 5 individuals using a DNeasy Blood & Tissue kit (Qiagen, Hilden, Germany). Extracted DNA was randomly fragmented to a size of 350bp then end-polished, A-tailed, and ligated with Illumina sequencing adapters using a NEBNext^®^ Ultra™ DNA Library Prep Kit (New England Biolabs, Ipswich MA, USA), and further PCR enriched with primers of P5 and P7 oligos. The PCR products which comprised the final libraries were purified (AMPure XP system, Beckman Coulter Life Sciences, Indianapolis IN, USA) and subjected to quality control tests that included size distribution by Agilent 2100 Bioanalyzer (Agilent Technologies, CA, USA), and molarity measurement using real-time PCR. The libraries were then pooled and sequenced on a NovaSeq 6000 (Illumina) using 2 × 150 bp chemistry by Novogene HK Company Limited, Hong Kong.

### Genome assembly

Quality filtering of raw sequencing reads was performed with Trimmomatic (Bolger *et al*., 2014), using the following settings: leading = 20, trailing = 20, slidingwindow = 4:20, minlen = 70. Adapter sequences were removed using the ILLUMINACLIP function, with maximum seed mismatches = 2 and the palindrome clip threshold = 30. Filtered reads were mapped to a *w*Mel reference genome (GenBank accession: NC_002978.6) and an *Ae. aegypti* mitochondrial reference genome (GenBank accession: NC_035159.1) using the Burrows-Wheeler Aligner (BWA), with the bwa mem algorithm and default parameters (Li, 2013). SAMtools and BCFtools were used to perform quality filtering of alignments and variant calling (Li *et al*., 2009; Danecek *et al*., 2021). PCR duplicates were excluded from downstream analyses by soft masking. Reads with a MAPQ score < 25 were removed from alignments, except for those with MAPQ = 0, which were retained to allow for mapping to repetitive regions. A maximum of 2000 reads per position were used to calculate genomic likelihoods. Ploidy was set to haploid for variant calling. The variant call output was used to create a consensus genome sequence for each sample, wherein low coverage positions (depth < 5) were masked as ‘N’. For *Wolbachia* genomes, Kraken2 (Wood *et al*., 2019) was used to search for sequence contamination with the Standard-8 precompiled reference database (https://benlangmead.github.io/aws-indexes/k2; downloaded 17/9/21). The sequencing reads mapped by bwa to the *w*Mel reference genome were filtered to remove reads matching taxa other than *Wolbachia*, and genome assemblies were then repeated with the filtered datasets, using the above pipeline. Genome sequences were inspected and aligned with Geneious v 9.1.8 (https://www.geneious.com). Read mapping densities were analysed with CNVpytor (Suvakov *et al*., 2021) using the read depth approach, in order to search for putative deletions and duplications within the *w*Mel genomes. Bin sizes of 100 and 500 were tested for the histogram, partition and CNV call steps.

### Phylogenetic analysis

A phylogenetic tree was constructed using the *w*Mel genomes from the present study and two large genome-wide SNP datasets that include representatives of the different *w*Mel clades, obtained from *D. melanogaster* sampled from various international locations (Richardson *et al*., 2012; Chrostek *et al*., 2013; Early and Clark, 2013). Polymorphic sites that were common to both datasets were combined into a single SNP matrix, with sites that contained ambiguous base calls in any sample excluded from the analysis. A total of 58 polymorphic loci were retained. Maximum likelihood trees were constructed with RAxML-HPC v8.2.12 (Stamatakis, 2014) on XSEDE, using a GTR-GAMMA model with the Lewis method of ascertainment bias correction (Lewis, 2001), 44 alignment patterns and rapid bootstrapping (100 inferences). Bootstrap scores were plotted onto the best scoring ML tree. RAxML-HPC was accessed through the CIPRES Science Gateway (Miller *et al*., 2010).

### Life history parameters and cytoplasmic incompatibility

To compare the effects of *w*MelM and *w*Mel infection on mosquito life history, we performed phenotypic assessments of the *w*Mel, *w*MelM and *w*Mel.tet populations. Fresh stored eggs (<1 week old) from each population were hatched in trays filled with 3L of RO water, a few grains of yeast, and one tablet of fish food. One day after hatching, 100 larvae were counted into trays filled with 500 mL of RO water and provided with fish food *ad libitum*, with 12 replicate trays per population. To determine the average larval development time for each tray, pupae were counted and sexed every day in the morning and evening. Pupae from each population were pooled across replicate trays (but with sexes separate) and used in the following experiments. The remaining pupae were pooled across sexes and released into BugDorm-1 cages for the quiescent egg viability experiment.

To determine adult longevity, 25 females and 25 males were placed in 3 L cages with cups of 10% sucrose and water, with eight replicate cages per population. Females were blood fed once per week. Dead mosquitoes were recorded and removed three times per week until all mosquitoes had died.

To determine female fertility and patterns of cytoplasmic incompatibility between *w*Mel variants, we set up reciprocal crosses between *w*Mel, *w*MelM and *w*Mel.tet adults. Fifty females (1 d old) from the *w*Mel, *w*MelM or *w*Mel.tet populations were aspirated into 3 L cages containing 50 *w*Mel, *w*MelM or *w*Mel.tet males (1 d old), for a total of 9 crosses. Five d old females were blood fed and 30 engorged females per cross were isolated for oviposition in 70 mL specimen cups with mesh lids that were filled with 15 mL of larval rearing water and lined with a strip of sandpaper (Norton Master Painters P80 sandpaper, Saint-Gobain Abrasives Pty. Ltd., Thomastown, Victoria, Australia). Eggs were collected, partially dried, and then hatched three days after collection. Female fecundity was determined by counting the total number of eggs on each sandpaper strip, while hatch proportions were determined by dividing the number of hatched eggs (with a clearly detached egg cap) by the total number of eggs per female. Females that did not lay eggs or died before laying eggs were excluded from the analysis.

To evaluate fertility across gonotrophic cycles, females from the within-strain crosses (e.g. *w*Mel x *w*Mel) were blood fed again after laying eggs. Eggs were collected from four gonotrophic cycles in total, when females were 9, 14, 19 and 23 d old.

As an estimate of body size, one wing each from 20 males and 20 females per population was dissected and measured for its length (Ross *et al*., 2016). Damaged wings excluded from the analysis.

### Quiescent egg viability

To test quiescent egg viability which is an important ecological trait that can be affected by *Wolbachia* infections (Lau *et al*., 2021), cages of 5 d old females from *w*Mel, *w*MelM or *w*Mel.tet populations were blood fed and six cups filled with larval rearing water and lined with sandpaper strips were placed inside in each cage. Eggs were collected five days after blood feeding, partially dried, then placed in a sealed chamber with an open container of saturated potassium chloride (KCl) solution to maintain a constant humidity of ~84%. When eggs were 1, 2, 4, 6, 8, 10, 12, 14, 16, 18, 20 and 22 weeks old, small sections of each sandpaper strip were removed and submerged in water with a few grains of yeast and fish food to hatch. Twelve replicate batches of eggs were hatched per population at each time point, with around 50-100 eggs per batch. Hatch proportions were determined by dividing the number of hatched eggs (with a clearly detached egg cap) by the total number of eggs in each batch.

### *Wolbachia* density across life stages

We compared the *Wolbachia* density of *w*MelM and *w*Mel during development by storing random subsets of 1^st^ instar larvae, 3^rd^ instar larvae, female pupae, male pupae, female adults and male adults (within 24 hr of emergence) in 100% ethanol. Fifteen individuals from each life stage, sex and *w*Mel variant were then measured for *Wolbachia* density using qPCR (see “*Wolbachia* detection and density”).

### DENV-2 virus challenge and quantification

We measured the vector competence of *w*MelM for DENV-2 relative to two other *Wolbachia* variants (*w*Mel and *w*AlbB) and a matched uninfected population (*w*Mel.tet). Experiments were performed in a quarantine insectary with *Ae. aegypti* held in incubators (PG50 Plant Growth Chambers, Labec Laboratory Equipment, Marrickville, NSW, Australia) set to a constant 26°C with a 12:12 light: dark photoperiod. Eggs (10 d old) from the *w*Mel, *w*MelM, *w*Mel.tet and *w*AlbB populations were hatched and larvae were reared to adulthood in five replicate trays of 100 larvae per temperature treatment for each population. Adults were released into 13.8-liter cages (BugDorm-4F2222, Megaview Science C Ltd., Taichung, Taiwan) and provided with 10% sucrose solution. Adults were starved for 24 hr before virus challenge and experiments were performed with 6-7d old females.

We performed virus challenge experiments with a 1 x 10^7^ TCID50u/mL dose of DENV-2 in human blood. DENV-2 (Cosmopolitan) provided by VDRL and isolated in Melbourne from a traveler in 2016 and was grown in C6/36 cells before use in experiments. Females were fed human blood sourced from the Red Cross under agreement number 16-10VIC-02 and spiked with DENV-2. Blood was provided through 6 mL Hemotek membrane feeders (Hemotek Ltd., Blackburn Lancashire, Great Britain) which were placed on the top of each cage for 30 min and heated with pocket hand warmers (Kathmandu). Engorged mosquitoes were transferred to cages with sucrose solution and cups lined with sandpaper strips for oviposition, with non-blood fed mosquitoes discarded. Females were maintained at 26°C for 12 d before processing. Cages were chilled briefly, then heads were dissected from bodies on a cold plate and stored individually in 1.7 mL tubes with 100 μL crushing media (DMEM with 1000u/ml streptomycin/penicillin and 2ug/ml amphoterin B) and two 3mm glass beads. Tubes were stored at - 20°C for 48 hr before removal from the quarantine insectary for long-term storage at −80°C.

Virus titres were quantified from whole bodies (heads removed) from 30 individuals per population with TCID_50_ assays as described previously (Duchemin *et al*., 2017). Briefly, 30 μl of crushed mosquito were serially 10-fold diluted in medium (DMEM containing 2% FBS). Using 96-well plates, 50 ul of each dilution was sequentially placed in wells (6 replicates). 100ul of fresh medium containing Vero cells (final cell confluency of 50-60%) was then overlaid. The cells were incubated for 7 days before they were observed for cytopathic effect (CPE). The 50% endpoint was calculated using the Reed and Muench method (Reed and Muench, 1938). The infection rate was calculated as the proportion (percentage) of all experimentally infected mosquitoes (n = 30) in which DENV was detected.

### Cytoplasmic incompatibility and *Wolbachia* density following cold and heat stress

We measured cytoplasmic incompatibility and *Wolbachia* density in adults after being exposed to cyclical cold or heat stress during the egg stage for one week. Eggs were collected on sandpaper from both *Wolbachia*-infected populations. Four days after collection, batches of 40-60 eggs were brushed off the sandpaper and tipped into 0.2 mL PCR tubes (12 replicate tubes per population) and exposed to cyclical temperatures of 3-13°C, 4-14°C, 6-16°C, 27-37°C, 28-38°C or 29-39°C for 7 d in Biometra TProfessional TRIO 48 thermocyclers (Biometra, Göttingen, Germany) according to Kong *et al*. (2016) and Ross *et al*. (2019b). Eggs of the same age from each population, as well as *w*Mel.tet eggs, were kept at 26°C. Eggs from all treatments were brought to 26°C and hatched synchronously. Larvae were reared at a controlled density (up to 100 larvae per tray of 500 mL water). Pupae were sexed and 8-30 adults per population were stored in absolute ethanol within 24 hr of emergence for *Wolbachia* density measurements (see *Wolbachia* detection and density). The remaining pupae were left to emerge into 3 L cages (with each sex, temperature treatment and population held in separate cages) for cytoplasmic incompatibility crosses. Due to low survivorship of eggs held at 3-13°C and 4-14°C, individuals from these treatments were not used in crossing experiments.

We established two sets of crosses to test cytoplasmic incompatibility induced by *Wolbachia*-infected males and test the ability of *Wolbachia*-infected females to restore compatibility with *Wolbachia*-infected males. In the first set, untreated *w*Mel.tet females were crossed with *Wolbachia*-infected males from each temperature treatment. In the second set, *Wolbachia*-infected females from each temperature treatment were crossed with untreated *Wolbachia*-infected males. Five-day old females were blood-fed and 20 females per cross were isolated for oviposition. Hatch proportions per female were determined according to previous experiments (see above).

### Maternal transmission of *Wolbachia*

We compared the ability of *w*Mel and *w*MelM females to transmit *Wolbachia* infections to their offspring after parental eggs were exposed to cyclical heat stress (28-38°C) for one week, or held at 26°C. Eggs were returned to 26°C and reared to adulthood, and *w*Mel and *w*MelM females from each temperature treatment were crossed to *w*Mel.tet males. Twenty females per cross were isolated for oviposition after blood feeding. We scored 10 offspring for their *Wolbachia* infection status from 10 females per infection type at each temperature to assess the loss of *Wolbachia* in next generation.

### *Wolbachia* detection and density

We used LAMP assays for rapid detection of *w*Mel during microinjection experiments according to Jasper *et al*. (2019) using *w*Mel-specific primer sets (Goncalves *et al*., 2019). qPCR assays were used to confirm the presence or absence of *Wolbachia* infection and measure relative density in all other experiments (Lee *et al*., 2012; Axford *et al*., 2016). DNA was extracted using 250 μL of 5% Chelex 100 resin (Bio-Rad oratories, Hercules, CA) according to methods described previously (Hoffmann *et al*., 2014). used a LightCycler^®^ 480 High Resolution Melting Master (HRMM) kit (Roche; Cat. No. 04909631001, Roche Diagnostics Australia Pty. Ltd., Castle Hill New South Wales, Australia) and IMMOLASETM DNA polymerase (5 U/μl) (Bioline; Cat. No. BIO-21047) as described by Lee *et al*. (2012). Three primer sets were used to amplify markers specific to mosquitoes (mRpS6_F 5’AGTTGAACGTATCGTTTCCCGCTAC3’ and mRpS6_R 5’ GAAGTGACGCAGCTTGTGGTCGTCC3’), *Ae. aegypti* (aRpS6_F 5’ATCAAGAAGCGCCGTGTCG3’ and aRpS6_R 5’CAGGTGCAGGATCTTCATGTATTCG3’), and *w*Mel (w1_F 5’AAAATCTTTGTGAAGAGGTGATCTGC3’ and w1_R 5’ GCACTGGGATGACAGGAAAAGG3’). Relative *Wolbachia* densities were determined by subtracting the Cp value of the *Wolbachia*-specific marker from the Cp value of the mosquito-specific marker. Differences in Cp were averaged across 2-3 consistent replicate runs, then transformed by 2^n^.

### Statistical analysis

All analyses were conducted using SPSS statistics version 26.0 for Mac (SPSS Inc, Chicago, IL). Egg hatch proportions were characterized per female (i.e. individual eggs were not treated as independent data points) and this trait as well as longevity were not normally distributed according to Shapiro-Wilk tests, therefore we analyzed these data with nonparametric log-rank tests for longevity and Kruskal-Wallis for egg hatch proportions. The development time for females and males, fecundity, *Wolbachia* density under heat and cold were analyzed by two-way ANOVAs. *Wolbachia* density for different life stages was compared with a t test. DENV-2 TCID_50_ comparisons across strains were undertaken using a Mann-Whitney test and differences in the infection rate were evaluated using two-tailed Fisher’s tests.

## Results

### Genomic analyses of the *w*MelM transinfection

The genome of the original *w*Mel transinfection (Walker *et al*., 2011) was very similar to both the *w*Mel reference genome (Wu *et al*., 2004) and the *w*Mel genome reported by Chrostek *et al*. (2013), differing from each by five and four polymorphic loci respectively, consistent with the three strains having originated from same *Drosophila* stock (*yw^67C23^*) less than 5,000 generations ago (Table S2). Phylogenetic analysis placed these genomes in a single monophyletic cluster together with the other members of *w*Mel clade III (Figure 1).

**Figure 1:**
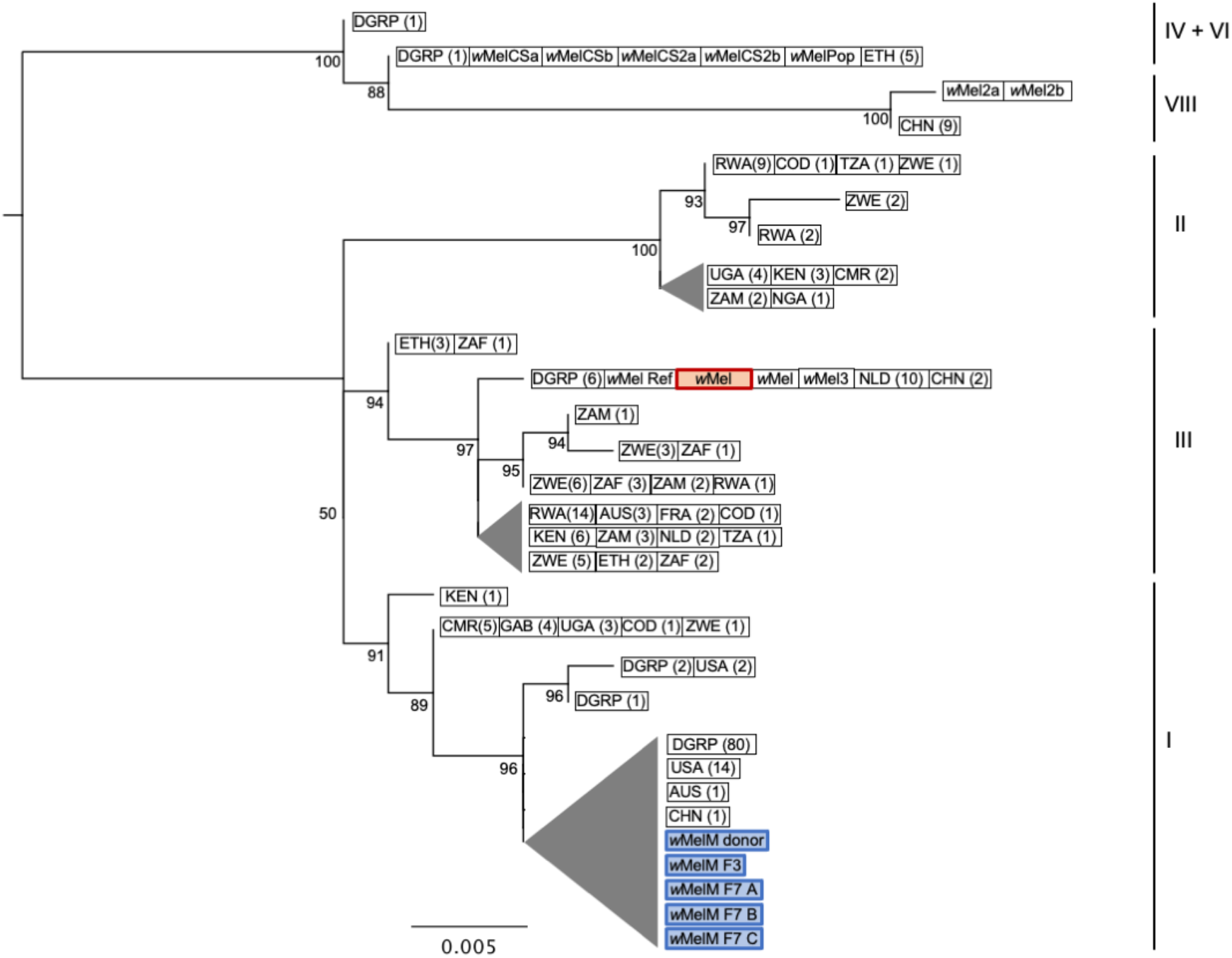
Genome-wide SNP phylogenetic analysis of *w*Mel variants. SNP data from the six genomes in this study and 253 previously published *w*Mel genomes were analysed (Richardson *et al*., 2012; Chrostek *et al*., 2013; Early and Clark, 2013). The *w*Mel variant highlighted in red is the original *w*Mel transinfection (Walker *et al*., 2011) used in phenotypic comparisons with *w*MelM. *w*MelM sequences are highlighted in blue. Maximum likelihood trees were constructed with RAxML-HPC using 58 SNP loci and ascertainment bias correction; scale bar = number of substitutions per SNP matrix site. Nodes with bootstrap values less than 50% have been collapsed; triangle height = length of longest branch within node. *Wolbachia w*Mel clades are shown on the right. In cases where multiple samples had identical SNP haplotypes, one representative sequence was used for tree construction. The number of sequences corresponding to each entry is shown in parentheses. For wild populations, the location of sampling is shown (refer to Table S1 for sample names). AUS = Australia; CHN = China; CMR = Cameroon; COD = Democratic Republic of the Congo; ETH = Ethiopia; FRA = France; GAB = Gabon; GIN = Guinea; KEN = Kenya; NGA = Nigeria; NLD = Netherlands; RWA = Rwanda; TZA = Tanzania; UGA = Uganda; USA = USA; ZAF = South Africa; ZMB = Zambia; ZWE = Zimbabwe; DGRP = Drosophila Genetic Reference Panel (originally sampled from Raleigh NC, USA).

Clear differences were observed between the genome sequence of the original *w*Mel transinfection strain and the *w*MelM variant. Phylogenetic analysis placed *w*MelM within *w*Mel clade I, with greatest similarity to *w*Mel variants from Ithaca NY, USA and Tasmania, Australia (Figure 1). Thirty-six SNPs and small indels were identified, with approximately half predicted to cause a change to an amino acid sequence (Table S2). Several of these changes were located within proteins with functions including transportation and secretion; amino acid, porphyrin, and nucleotide metabolism; and DNA replication and repair, while many others were located within hypothetical proteins of unknown function. Most SNPs identified were the same SNPs identified by Hague *et al*. (2022) in their comparison of *w*Mel genomes from temperate (clade III) and tropical (clade I) *D. melanogaster*, with five of these being significantly associated with temperature according to their analysis (P < 0.05 see Hague *et al*., 2022). Analysis of read mapping densities, performed with CNVpytor, did not identify any regions of significant copy number variation within the *w*MelM genome, when compared to the *w*Mel reference genome. We did, however, identify two large deletions through standard genome assembly – one within a gene encoding a S49 family peptidase and one within an ankyrin repeat containing gene.

Examination of allele frequencies indicated that in addition to *w*MelM, a *w*Mel variant with high identity to the *w*Mel reference sequence is likely to have been present within the *Drosophila w*MelM donor population, but only *w*MelM appears to have been retained within the transinfected mosquito populations. The *w*MelM genome sequences in *Ae. aegypti* at F_3_ and three isofemale lines at F_7_ were identical, suggesting no genetic changes across several generations following transinfection. Relative to the *Ae. aegypti w*MelM genomes, the *Drosophila w*MelM consensus sequence contained eight SNPs and lacked the two large deletions mentioned above. It is probable that these differences are artifacts introduced by the presence of the other *w*Mel variant in this sample.

The mitochondrial genome sequences of the five mosquito populations included in the study were very similar, with only five SNPs observed between them, four of which were located outside of known coding regions. Most of the SNPs were associated with anomalously low read depths, relative to their surrounding positions, making it likely that they represent sequencing artifacts.

### Limited effects of *w*MelM on life history traits

We compared the effects of *w*Mel and *w*MelM infection on *Ae. aegypti* life history traits. Overall, *w*MelM infection had no effect, or slightly increased mosquito fitness compared to uninfected (*w*Mel.tet) mosquitoes (Figure 2). *Wolbachia* infection had no clear effect on development time (two-way ANOVA: P = 0.124 for females and 0.194 for males, Figure 2A and 2B). Both *w*Mel variants increased adult female longevity (Log-rank test: *w*Mel vs *w*Mel.tet, P = 0.011, χ^2^ = 6.436, df = 1, *w*MelM vs *w*Mel.tet: P = 0.019, χ^2^ = 5.501, df = 1, Figure 2C) relative to *w*Mel.tet, but male longevity was unaffected by *Wolbachia* infection status (Log-rank test: *w*Mel vs *w*Mel.tet, P = 0.251, χ^2^ = 1.32, df = 1, *w*MelM vs *w*Mel.tet: P = 0.091, χ^2^ = 2.849, df = 1, Figure 2D). Fecundity was influenced by *Wolbachia* infection type (two-way ANOVA: F_2,326_ = 3.753, P = 0.024), with *w*MelM females consistently laying more eggs than the *w*Mel and *w*Mel.tet females (Figure 2E). Median egg hatch proportions were close to 100% across all gonotrophic cycles, regardless of *Wolbachia* infection type (Figure 2F). *w*Mel males had smaller wings than *w*Mel.tet males (one-way ANOVA: P = 0.04, Figure 1H) with no difference between the two *w*Mel variants (P = 0.767), while female wing length was unaffected by *Wolbachia* infection type (P>0.05, Figure 2G).

**Figure 2.**
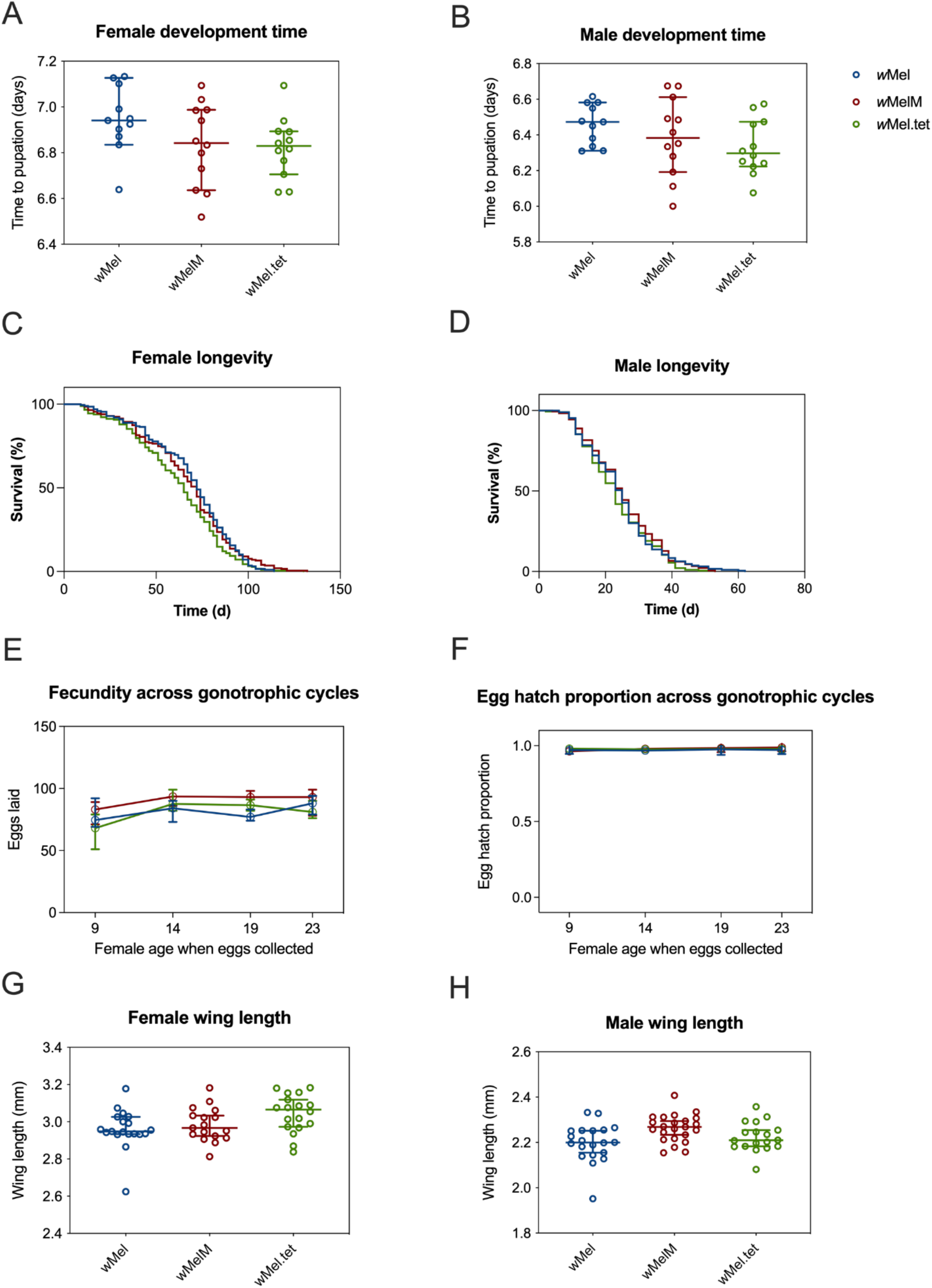
Life history parameters of backcrossed *w*Mel, *w*MelM and *w*Mel.tet *Aedes aegypti* populations. The *w*Mel (blue), *w*MelM (red) and *w*Mel.tet (green) populations were evaluated for the following traits: larval development time for (A) females and (B) males, longevity of (C) females and (D) males, (E) female fecundity across gonotrophic cycles, (F) egg hatch proportion across gonotrophic cycles, and wing length of (G) females and (H) males. Each point represents data averaged across a replicate container of 100 individuals (A-B) or data from individual mosquitoes (panels G-H). Medians and 95% confidence intervals are shown in lines and error bars.

### *w*Mel and *w*MelM induce complete cytoplasmic incompatibility

We tested the ability of *w*Mel and *w*MelM-infected males to induce cytoplasmic incompatibility with *w*Mel.tet females. These females produced no viable offspring when crossed to *w*Mel and *w*MelM males, indicating that both variants induce complete cytoplasmic incompatibility (Table 1). Reciprocal crosses between *w*Mel and *w*MelM variants resulted in high egg hatch proportions (>90%), indicating that the two variants are bidirectionally compatible (Table 1).

**Table 1.** Cytoplasmic incompatibility between *w*Mel, *w*MelM and *w*Mel.tet *Aedes aegypti* populations.

|  |  | Female |  |  |
| --- | --- | --- | --- | --- |
|  |  | wMel.tet | wMel | wMelM |
| Male | wMel.tet | 0.980 (0.967, 1) | 0.930 (0.867, 0.969) | 0.952 (0.904, 0.990) |
|  | wMel | 0 (0, 0) | 0.971 (0.945, 0.983) | 0.981 (0.955, 0.990) |
|  | wMelM | 0 (0, 0) | 0.963 (0.850, 0.986) | 0.962 (0.942, 0.986) |

Median egg hatch proportions are shown with 95% confidence intervals (lower, upper).

### *w*Mel variants decrease quiescent egg viability

Stored eggs from each population were hatched every two weeks to determine quiescent egg viability. Egg viability decreased across time for all three infection types, but hatch proportions for both *w*Mel variants were lower than *w*Mel.tet from week 2 onwards (Kruskal-Wallis test, week 2: H = 12.434, df = 2, *P* = 0.002, Figure 3). *w*Mel and *w*MelM did not differ significantly in hatch proportion across all weeks combined (statistical test *P* = 0.114), suggesting that the two variants have similar quiescent egg viability.

**Figure 3.**
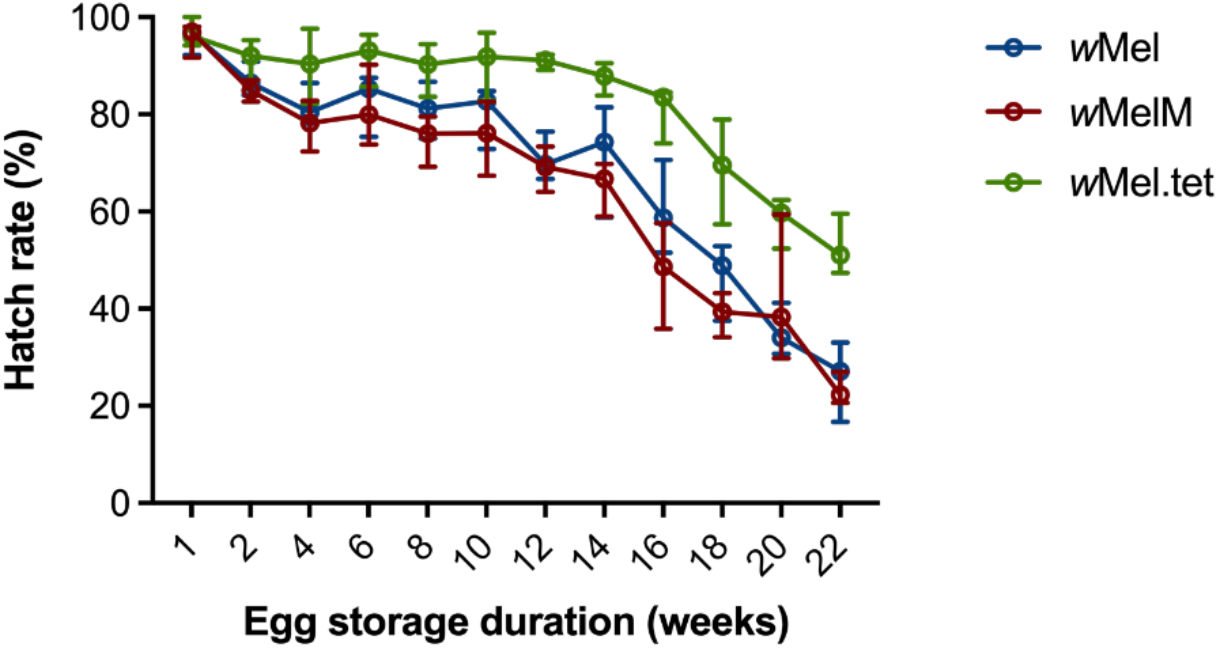
Quiescent egg viability of backcrossed *w*Mel, *w*MelM and *w*Mel.tet *Aedes aegypti* populations across 22 weeks. Symbols show median egg hatch proportions while error bars show 95% confidence intervals.

### *w*Mel variants have similar *Wolbachia* density across life stages

*Wolbachia* density in whole individuals differed across life stages (two-way ANOVA: F_5,177_ = 75.85, *P* < 0.001), with low densities observed in third instar larvae (Figure 4). We did not find differences in pairwise comparisons between the *w*Mel and *w*MelM strains (t-test: all *P* > 0.05) except for male pupae (*P* = 0.011), where *w*MelM male pupae had higher *Wolbachia* densities than *w*Mel male pupae (Figure 4). Overall, the *Wolbachia* density of *w*MelM was slightly higher than *w*Mel across all life stages but this effect was not significant (two-way ANOVA: F_1,177_ = 2.636, *P* = 0.106).

**Figure 4.**
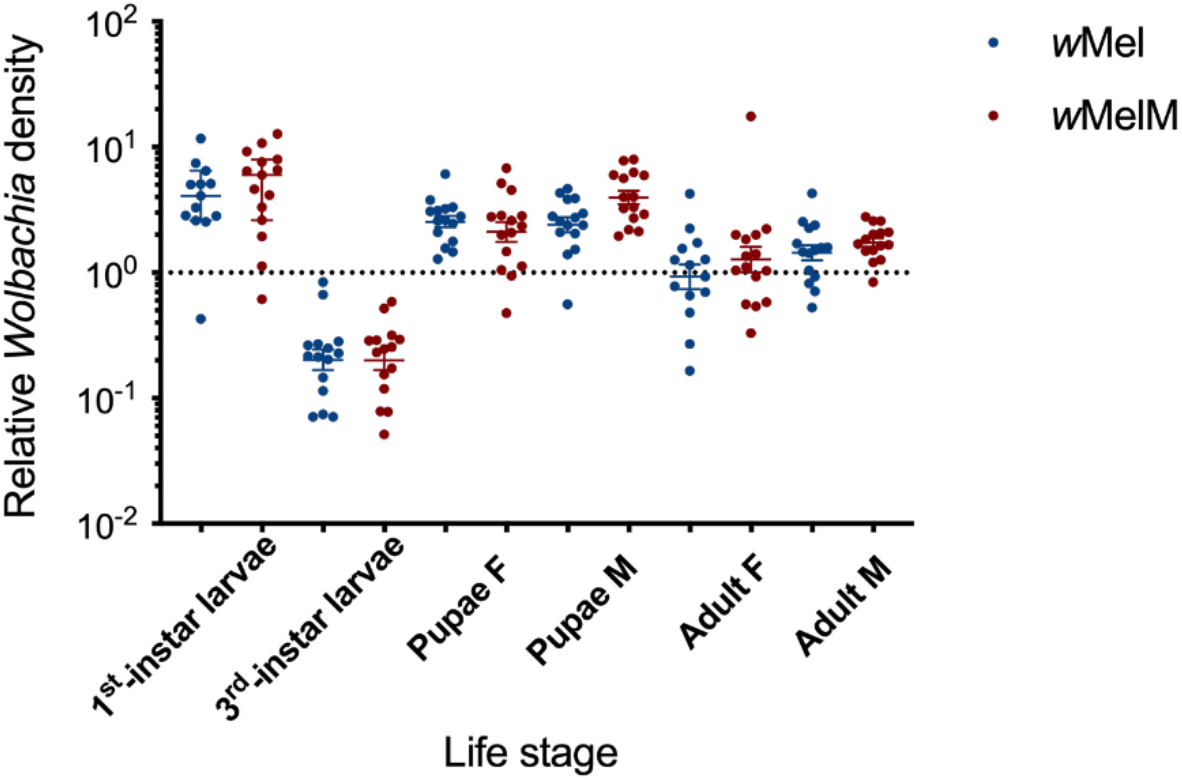
*Wolbachia* density of different life stages in backcrossed *w*Mel and *w*MelM *Aedes aegypti* populations. Each point represents the relative density for an individual averaged across 2-3 technical replicates. Medians and 95% confidence intervals are shown in lines and error bars. “F” stands for females and “M” stands for males.

### *w*MelM blocks DENV-2

We assessed the vector competence of *w*MelM-infected *Ae. aegypti* by feeding mosquitoes an infectious blood meal and comparing DENV-2 titres against *w*Mel, *w*AlbB and *w*Mel.tet populations. DENV-2 titres (*P* = 0.016) and the proportion of DENV-2 infected females (*P*=0.033) were significantly lower in *w*MelM compared to *w*Mel.tet (Figure 5). There was no detectable difference between *w*MelM blockage and that of the other two *Wolbachia* strains, *w*Mel and *w*AlbB, as measured by both DENV-2 replication and infection proportion (all *P* > 0.05).

**Figure 5.**
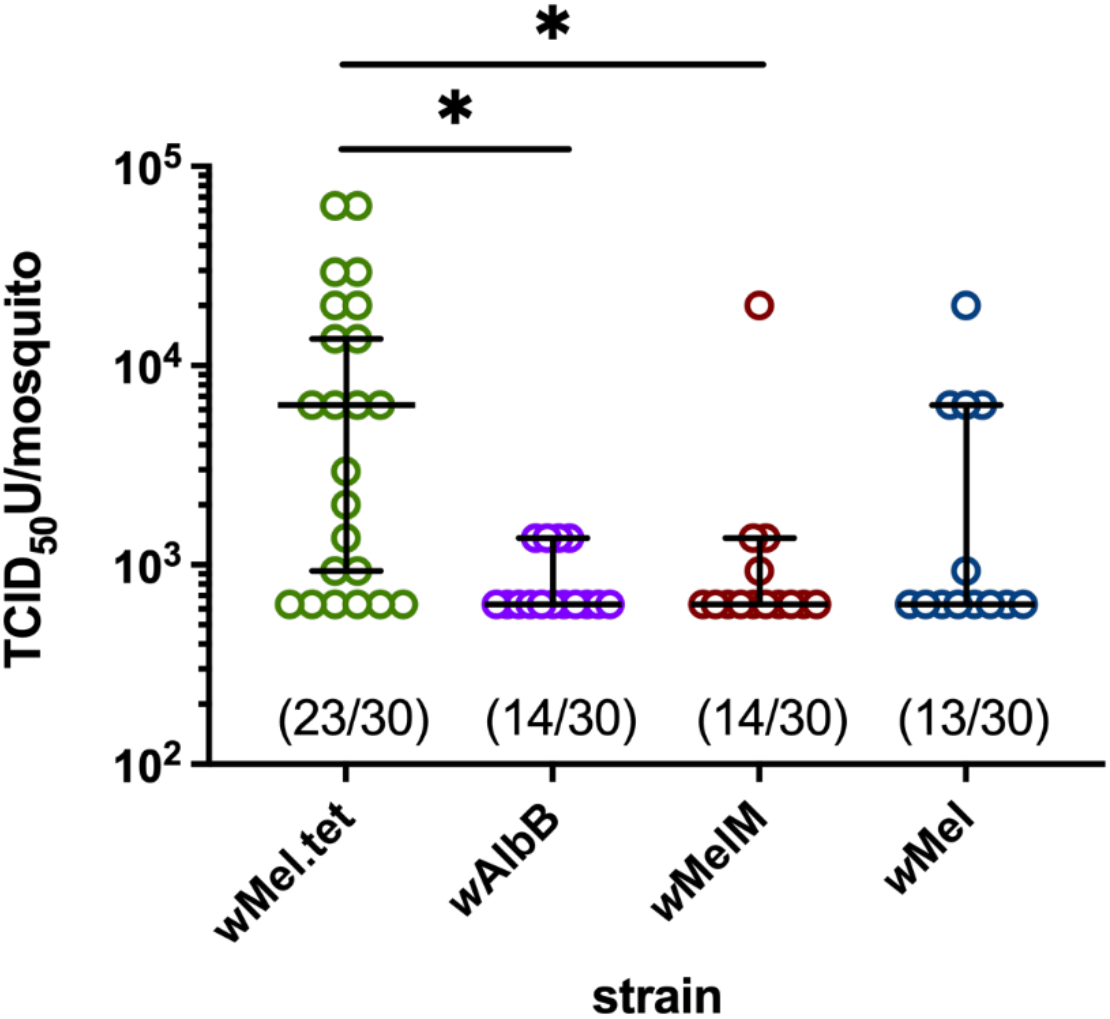
*w*MelM inhibits DENV-2 replication in virus-fed *Ae*. *aegypti*. Statistical comparisons of DENV-2 titre were compared using a Mann-Whitney test where * indicates *P* < 0.05. Each point on the plot represents an individual infected mosquito (with negative mosquitoes excluded), and lines and error bars denote medians ± 95% CIs. The proportion of DENV-2 positive mosquitoes out of the total tested (proportion infected) is indicated below each plot.

### *w*MelM induces stronger cytoplasmic incompatibility than *w*Mel under cyclical heat stress

We tested the ability of *w*Mel and *w*MelM-infected males to induce cytoplasmic incompatibility when eggs were exposed to cold or heat stress. Males from both *w*Mel and *w*MelM lines induced complete cytoplasmic incompatibility with *w*Mel.tet females at 6-16°C and 26°C (Figure 6A). Incomplete cytoplasmic incompatibility was observed at 27-37°C and above, with an increasing proportion of viable offspring produced as the temperature increased (Figure 6A). However, *w*MelM retained stronger cytoplasmic incompatibility than *w*Mel at high temperatures, with significantly lower egg hatch proportions when males were exposed to 28-38°C (Kruskal-Wallis test: H = 6.743, df = 1, *P* = 0.009) and 29-39°C (H = 5.344, df = 1, *P* = 0.021).

**Figure 6.**
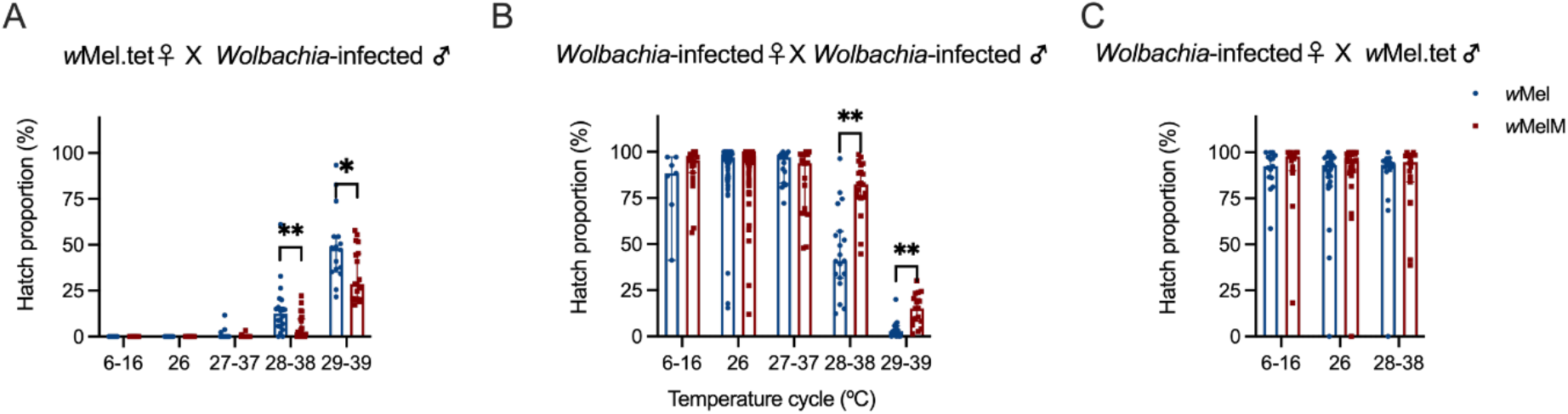
Cytoplasmic incompatibility induction and compatibility restoration by *w*Mel/*w*MelM variants under cold and heat stress. We performed crosses to test the ability of (A) *Wolbachia*-infected males to induce cytoplasmic incompatibility with *w*Mel.tet females and (B) *Wolbachia*-infected females to restore compatibility with *Wolbachia*-infected males. We also evaluated egg hatch proportions from crosses between (C) *Wolbachia*-infected females and *w*Mel.tet males used in the maternal transmission experiment. In each cross, males (A) or females (B, C) were exposed to cyclical temperatures for 7 d or held at 26°C during the egg stage. Each point represents the hatch proportion of eggs from a single female. Bars show medians while error bars show 95% confidence intervals.

*Wolbachia*-infected females exposed to cyclical temperatures as eggs were crossed to *Wolbachia*-infected males reared at 26°C to test the ability of females to restore compatibility (Figure 6B). Compared to *w*MelM, *w*Mel had a slightly decreased hatch rate under cold stress but treatments did not differ significantly (H = 1.838, df = 1, *P* = 0.175). After the exposure to cyclical temperatures of 27-37°C for 7 d, hatch rate reduced. Hatch rates in both strains decreased with temperature, but wMelM had a higher hatch rate under cyclical temperatures of 28-38°C (Kruskal-Wallis test: H=18.000, df=1, *P* < 0.001) and 29-39°C (H=13.766, df=1, *P* < 0.001).

### wMelM has increased maternal transmission fidelity compared to *w*Mel under cyclical heat stress

We compared the ability of *w*Mel and *w*MelM females to transmit *Wolbachia* to their offspring when they were exposed to 28-38°C as eggs for 7d or reared at a constant 26°C (Figure 6C). *w*Mel and *w*MelM females transmitted the infection to all their offspring at 26°C. In contrast, *w*Mel and *w*MelM-infected females failed to transmit the infection to some of their progeny at 28-38°C (Figure 7). More progeny from *w*MelM mothers had *Wolbachia* than from *w*Mel mothers (66.7% ± 6.87%) at a cycling 28-38°C (Kruskal-Wallis test: H=3.891, df=1, *P*= 0.048, Figure 7), suggesting that the transmission of *w*MelM may be more stable at high temperatures.

**Figure 7.**
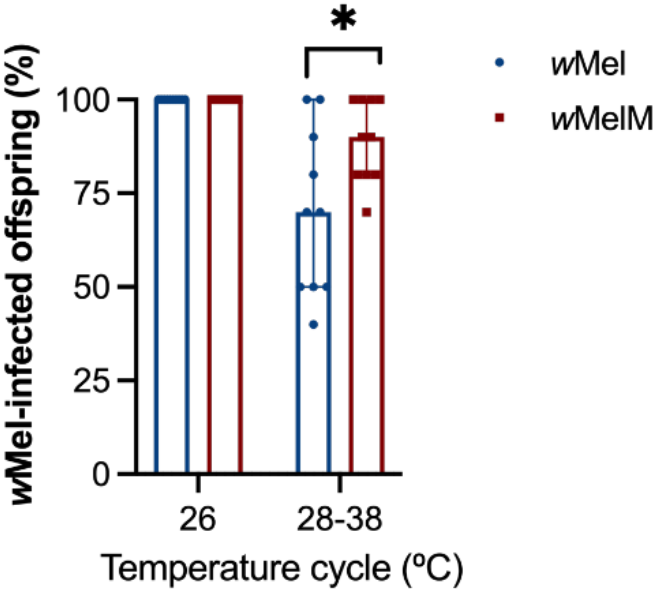
Maternal transmission of *w*Mel/*w*MelM variants following maternal heat stress exposure. *Wolbachia*-infected females were exposed to cyclical temperatures of 28-38°C for 7 d or held at 26°C during the egg stage, then crossed to *w*Mel.tet males. Each point represents the percent of *Wolbachia*-positive offspring from a single female. Medians and 95% confidence intervals are shown in lines.

### *w*MelM has a higher density than *w*Mel under heat and cold stress

We measured *Wolbachia* density when eggs were exposed to cold or heat stress. We found no significant effect of sex on *Wolbachia* density (*P* > 0.05), so data from males and females were pooled for the following analysis. In the two-way ANOVA, *Wolbachia* density was influenced both by the *w*Mel variant (F_1,142_ = 26.425, *P* < 0.001) and temperature (F_2,142_ = 247.264, *P* < 0.001). At high temperatures, the density of *w*Mel and *w*MelM decreased sharply (Figure 8B). For the lower temperature, the *Wolbachia* density was relatively stable but was still influenced by *w*Mel variant (F_1,99_ = 12.658, *P* = 0.001) and temperature (F_2,99_ = 18.323, *P* < 0.001) (Figure 8A). Although *w*Mel and *w*MelM had similar density at the adult stage when reared at constant 26°C (Figure 4), we did find that *w*MelM had a higher *Wolbachia* density under all the cycling heat and cold stress regimes (Figure 8). Compared to *w*Mel, the density of *w*MelM was around 2 times higher overall in this experiment.

**Figure 8.**
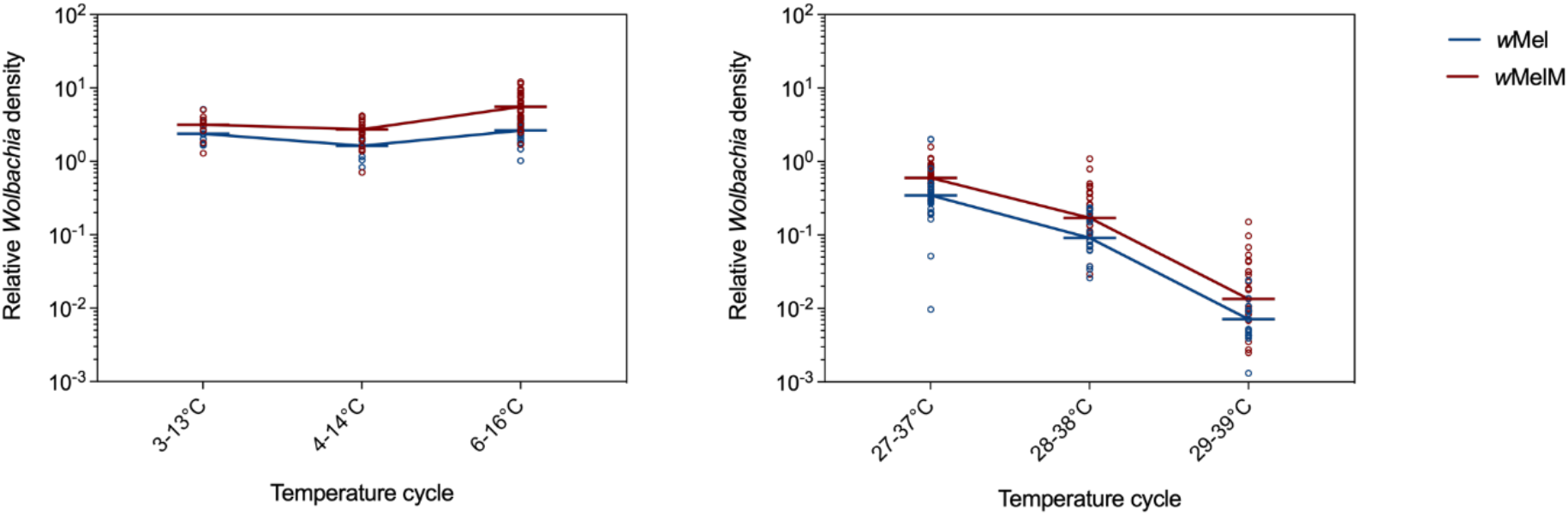
*Wolbachia* density in adult mosquitoes following exposure to cyclical (A) cold and (B) heat stress during the egg stage. Eggs were exposed to cyclical temperatures for 7 d. Each point represents the relative density for an individual averaged across 2-3 technical replicates. Medians are shown as short horizontal lines and lines join medians across different temperature cycles.

## Discussion

We successfully used microinjection to transfer *Wolbachia* directly from *D. melanogaster* to *Ae. aegypti* to generate the *w*MelM line. *w*MelM differs from the original *w*Mel transinfection (Walker *et al*., 2011) both in terms of the source of the *w*Mel material (laboratory versus field-derived *D. melanogaster*) and initial passage through a cell line. Although different methods were used to develop the *w*Mel and *w*MelM transinfection, the *w*MelM variant like the original *w*Mel has minimal effects on mosquito fitness. The *w*MelM variant was associated with a somewhat higher host fecundity compared to the *w*Mel strain and a longer host life span when compared to *w*Mel.tet mosquitoes, traits that would be likely to improve the success of release strategies in a field setting aimed at replacement. Apart from these fitness effects, we found minor differences in the density of *Wolbachia* which may be correlated with the degree of virus blocking (Lu *et al*., 2012; Osborne *et al*., 2012; Fraser *et al*., 2017). For *w*MelM, *Wolbachia* density was higher but only under cycling heat and cold conditions; this leads to the expectation that virus blockage should be similar in the two strains, consistent with the similar virus blocking ability for *w*MelM, *w*Mel and *w*AlbB observed at 26°C. The high level of transmission and incompatibility generated by *w*MelM under the varied thermal conditions tested here may facilitate its invasion into natural field populations and persistence at a high level.

In native *Drosophila* hosts, *w*Mel *Wolbachia* density can vary substantially (Early and Clark, 2013); for instance, individuals from Ithaca NY, USA, most of which were infected with *w*Mel from clade I, had higher *Wolbachia* titres when reared at room temperature than individuals from Tasmania, Australia, Zimbabwe and the Netherlands, most of which were infected with *w*Mel from clade III. It is possible that a common mechanism underlies this finding and our observation of higher densities of *w*MelM (clade I) in temperature-stressed *Ae. aegypti* relative to the original *w*Mel transinfection strain (clade III). However, the cause of these differences in *Wolbachia* density remain to be determined. In more distantly related *w*MelCS-like variants (clade VI), high *Wolbachia* titres are associated with both the loss of and the amplification of the Octomom genome region (Chrostek *et al*., 2013; Chrostek and Teixeira, 2015; Duarte *et al*., 2021). In our case, we did not detect any alteration of Octomom copy number in *w*MelM. However, we did observe non-silent changes in genes encoding a *Wolbachia* surface protein (*wspB*), a member of the type IV secretion system, and proteins involved in amino acid metabolism, all functions that are likely to be important in interactions between endosymbiont and host (Pichon *et al*., 2009; Caragata *et al*., 2014; Rice *et al*., 2017; Jimenez *et al*., 2019; Epis *et al*., 2020).

Previous studies have shown that strong geographic structuring exists among global *w*Mel populations. While multiple *w*Mel variants can often be found within a single location or region, usually one variant predominates at any given location, suggestive of adaptation to local environmental conditions (Richardson *et al*., 2012; Early and Clark, 2013). There have been very few changes in the original *w*Mel strain since transinfection (Huang *et al*., 2020) but this strain is different genomically from some *w*Mel strains present in *D. melanogaster* from Australia (which provided the donor for *w*MelM). Phylogenetic analysis placed the original *w*Mel transinfection (Walker *et al*., 2011) within clade III, which contains 13 of the 14 other Australian (Tasmanian) samples included in the tree, and *w*MelM within clade I, which contains a single Tasmanian sample. This pattern raises the possibility that clade III *w*Mel variants may have a competitive advantage over clade I variants in some parts of southern Australia, such as Tasmania, which lies at the southern boundary of the cline of decreasing *Wolbachia* infection incidence (Kriesner *et al*., 2016). Most of the SNPs in the *w*MelM genome were identified in a study of *w*Mel SNPs that distinguished different geographic climate zones, with five of these shown to be significantly associated with temperature (Hague *et al*., 2022). In our analysis, we did find a SNP in the gene encoding the *Wolbachia* outer membrane protein B (*wspB*, WD0009), which results in a premature stop codon and is likely a major determinant of *Wolbachia* thermal sensitivity (Hague *et al*., 2022). It may be that the polymorphisms in the *w*MelM genome and the higher densities observed for *w*MelM under heat and cold stress are selected against in regions with lower yearly minimum temperatures (Kriesner *et al*., 2016). It would be interesting to determine the frequency of different *w*Mel variants in natural *D. melanogaster* populations across Australia, especially in more northerly regions where *Ae. aegypti* population replacement programs are being conducted.

Phenotypic differences between *w*Mel and *w*MelM may relate to the donor line of natural *Drosophila* used in the transinfection as well as the difference in methodology related to mosquito cell passaging rather than direct transfer. In eastern Australian *D. melanogaster, Wolbachia* were first identified from incompatibility generated through crossing experiments between populations from along a latitudinal cline (Hoffmann, 1988) and *w*Mel shows a stable latitudinal pattern from being near 100% in incidence to being at a very low frequency (Kriesner *et al*., 2016). This pattern suggests environmental effects on *Wolbachia* dynamics and/or differences in the *Wolbachia* strain/host backgrounds affecting these dynamics. Transplant experiments followed by semi-field cage experiments point to a complex pattern of *Wolbachia* and host associated fitness effects at ends of the cline (Olsen *et al*., 2001) while experimental studies implicate environmental effects interacting with *Wolbachia* fitness (Kriesner *et al*., 2016). Other data also point to different *w*Mel variants changing over time at different temperatures (Versace *et al*., 2014). Furthermore, environmental and genetic difference sometimes contribute together to *Wolbachia* transmission and frequency variation expected to influence the *Wolbachia* spread (Hague *et al*., 2020b).

For *Wolbachia* infections generally, high temperatures can have an impact on release success by weakening the reproductive effects induced by *Wolbachia* (Trpis *et al*., 1981; Johanowicz and Hoy, 1998; Ross *et al*., 2017) and *Wolbachia* can even be eliminated under sufficient thermal stress (Trpis *et al*., 1981; Stouthamer *et al*., 1990). In the case of *Ae. aegypti*, high temperature effects depend on the *Wolbachia* strain, with *w*AlbB performing better than *w*Mel (Ross *et al*., 2017). The higher *Wolbachia* density of *w*AlbB under heat may explain the fact that its phenotypic effects are not altered much by high temperatures. This also appears to be the case for the *w*MelM variant.

Given the impact of high temperature on the dynamics of *Wolbachia*-infected mosquitoes and the benefits of *Wolbachia* releases targeting *Ae. aegypti* in several countries from different climate zones where dengue is common (Bhatt *et al*., 2013; Ritchie, 2018), it may be prudent to consider multiple strains in releases in different environments (Ross *et al*., 2019a). The *w*MelM variant generated here has minimal fitness effects but a higher phenotypic stability at high temperatures compared to *w*Mel. Our results suggest that this new transinfected strain may be useful in environments where temperatures are variable. Given that temperature conditions also change seasonally, the new strain may allow for releases in spring when temperatures are often variable. It is also worth considering other strains from field populations for transfections.

## Acknowledgements

We thank Kelly Richardson for collecting the *Drosophila* line used for microinjection, Marianne P. Coquilleau for the assistance in the experiments and Moshe Jasper for support with LAMP assays.

## Funding

AAH was supported by the National Health and Medical Research Council (1132412, 1118640, www.nhmrc.gov.au). PAR was supported by a University of Melbourne Early Career Research Grant. The funders had no role in study design, data collection and analysis, decision to publish, or preparation of the manuscript.

## Data and strain availability

The *w*MelM *Aedes aegypti* strain is available from the authors on request.

## Notes

### Competing Interest Statement

The authors have declared no competing interest.

## References

Ahmad, N.A., Mancini, M.V., Ant, T.H., Martinez, J., Kamarul, G.M.R., Nazni, W.A. et al. (2021) *Wolbachia* strain *w*AlbB maintains high density and dengue inhibition following introduction into a field population of *Aedes aegypti*. Philos Trans R Soc Lond B Biol Sci 376: 20190809.

Axford, J.K., Ross, P.A., Yeap, H.L., Callahan, A.G., and Hoffmann, A.A. (2016) Fitness of *w*AlbB *Wolbachia* infection in *Aedes aegypti*: parameter estimates in an outcrossed background and potential for population invasion. Am J Trop Med Hyg 94: 507–516.

Beebe, N.W., Pagendam, D., Trewin, B.J., Boomer, A., Bradford, M., Ford, A. et al. (2021) Releasing incompatible males drives strong suppression across populations of wild and *Wolbachia*-carrying *Aedes aegypti* in Australia. P Natl Acad Sci USA 118.

Bhatt, S., Gething, P.W., Brady, O.J., Messina, J.P., Farlow, A.W., Moyes, C.L. et al. (2013) The global distribution and burden of dengue. Nature 496: 504–507.

Bolger, A.M., Lohse, M., and Usadel, B. (2014) Trimmomatic: a flexible trimmer for Illumina sequence data. Bioinformatics 30: 2114–2120.

Bull, J.J., and Turelli, M. (2013) *Wolbachia* versus dengue: evolutionary forecasts. Evol Med Public Health 2013: 197–207.

Burdina, E.V., Bykov, R.A., Menshanov, P.N., Ilinsky, Y.Y., and Gruntenko, N.E. (2021) Unique *Wolbachia* strain *w*MelPlus increases heat stress resistance in *Drosophila melanogaster*. Arch Insect Biochem 106: e21776.

Caragata, E.P., Rances, E., O’Neill, S.L., and McGraw, E.A. (2014) Competition for amino acids between *Wolbachia* and the mosquito host, *Aedes aegypti*. Microb Ecol 67: 205–218.

Caspari, E., and Watson, G.S. (1959) On the evolutionary importance of cytoplasmic sterility in mosquitoes. Evolution 13: 568–570.

Chrostek, E., and Teixeira, L. (2015) Mutualism breakdown by amplification of *Wolbachia* genes. PLoS Biol 13: e1002065.

Chrostek, E., Martins, N.E., Marialva, M.S., and Teixeira, L. (2021) *Wolbachia*-conferred antiviral protection Is determined by developmental temperature. bioRxiv 2020-06.

Chrostek, E., Marialva, M.S.P., Esteves, S.S., Weinert, L.A., Martinez, J., Jiggins, F.M., and Teixeira, L. (2013) *Wolbachia* variants induce differential protection to viruses in *Drosophila melanogaster:* a phenotypic and phylogenomic analysis. PLoS Genet 9: e1003896.

Corbin, C., Heyworth, E.R., Ferrari, J., and Hurst, G.D.D. (2017) Heritable symbionts in a world of varying temperature. Heredity 118: 10–20.

Crawford, J.E., Clarke, D.W., Criswell, V., Desnoyer, M., Cornel, D., Deegan, B. et al. (2020) Efficient production of male *Wolbachia*-infected *Aedes aegypti* mosquitoes enables large-scale suppression of wild populations. Nat Biotechnol 38: 482–492.

Danecek, P., Bonfield, J.K., Liddle, J., Marshall, J., Ohan, V., Pollard, M.O. et al. (2021) Twelve years of SAMtools and BCFtools. Gigascience 10: giab008.

Duarte, E.H., Carvalho, A., Lopez-Madrigal, S., Costa, J., and Teixeira, L. (2021) Forward genetics in *Wolbachia:* regulation of *Wolbachia* proliferation by the amplification and deletion of an addictive genomic island. PLoS Genet 17: e1009612.

Duchemin, J.B., Mee, P.T., Lynch, S.E., Vedururu, R., Trinidad, L., and Paradkar, P. (2017) Zika vector transmission risk in temperate Australia: a vector competence study. Virol J 14: 1–10.

Early, A.M., and Clark, A.G. (2013) Monophyly of *Wolbachia* pipientis genomes within *Drosophila melanogaster:* geographic structuring, titre variation and host effects across five populations. Mol Ecol 22: 5765–5778.

Epis, S., Varotto-Boccazzi, I., Crotti, E., Damiani, C., Giovati, L., Mandrioli, M. et al. (2020) Chimeric symbionts expressing a *Wolbachia* protein stimulate mosquito immunity and inhibit filarial parasite development. Commun Biol 3: 1–10.

Fraser, J.E., De Bruyne, J.T., Iturbe-Ormaetxe, I., Stepnell, J., Burns, R.L., Flores, H.A., and O’Neill, S.L. (2017) Novel *Wolbachia*-transinfected *Aedes aegypti* mosquitoes possess diverse fitness and vector competence phenotypes. PLoS Pathog 13: e1006751.

Garcia, G.D., dos Santos, L.M.B., Villela, D.A.M., and Maciel-de-Freitas, R. (2016) Using *Wolbachia* releases to estimate *Aedes aegypti* (Diptera: Culicidae) population size and survival. PLoS One 11: e0160196.

Garcia, G.D., Sylvestre, G., Aguiar, R., da Costa, G.B., Martins, A.J., Lima, J.B.P. et al. (2019) Matching the genetics of released and local *Aedes aegypti* populations is critical to assure *Wolbachia* invasion. PLoS Neglect Trop D 13: e0007023.

Goncalves, D.D., Hooker, D.J., Dong, Y., Baran, N., Kyrylos, P., Iturbe-Ormaetxe, I. et al. (2019) Detecting *w*Mel *Wolbachia* in field-collected *Aedes aegypti* mosquitoes using loop-mediated isothermal amplification (LAMP). Parasite Vector 12: 1–5.

Gruntenko, N.E., Ilinsky, Y.Y., Adonyeva, N.V., Burdina, E.V., Bykov, R.A., Menshanov, P.N., and Rauschenbach, I.Y. (2017) Various *Wolbachia* genotypes differently influence host *Drosophila* dopamine metabolism and survival under heat stress conditions. Bmc Evol Biol 17: 15–22.

Guzman, M.G., Halstead, S.B., Artsob, H., Buchy, P., Jeremy, F., Gubler, D.J. et al. (2010) Dengue: a continuing global threat. Nat Rev Microbiol 8: S7–S16.

Hague, M.T., Shropshire, J.D., Caldwell, C.N., Statz, J.P., Stanek, K.A., Conner, W.R., and Cooper, B.S. (2022) Temperature effects on cellular host-microbe interactions explain continent-wide endosymbiont prevalence. Curr Biol 32: 1–11.

Hague, M.T.J., Mavengere, H., Matute, D.R., and Cooper, B.S. (2020) Environmental and genetic contributions to imperfect *w*Mel-Like *Wolbachia* transmission and frequency variation. Genetics 215: 1117–1132.

Hien, N.T., Anh, D.D., Le, N.H., Yen, N.T., Phong, T.V., Nam, V.S. et al. (2021) Environmental factors influence the local establishment of *Wolbachia* in *Aedes aegypti* mosquitoes in two small communities in central Vietnam. Gates Open Research 5: 147.

Hoffmann, A.A. (1988) Partial cytoplasmic incompatibility between two Australian populations of *Drosophila melanogaster*., 48(1), 61-67. Entomol Exp Appl 48: 61–67.

Hoffmann, A.A., and Turelli, M. (1997) Cytoplasmic incompatibility in insects In: O’Neill SL, Hoffmann AA, Werren JH, editors. Influential passengers: inherited microorganisms and arthropod reproduction:42–80.

Hoffmann, A.A., Turelli, M., and Simmons, G.M. (1986) Unidirectional incompatibility between populations of *Drosophila simulans*. Evolution 40: 692–701.

Hoffmann, A.A., Ross, P.A., and Rasic, G. (2015) *Wolbachia* strains for disease control: ecological and evolutionary considerations. Evol Appl 8: 751–768.

Hoffmann, A.A., Iturbe-Ormaetxe, I., Callahan, A.G., Phillips, B., Billington, K., Axford, J.K. et al. (2014) Stability of the *w*Mel *Wolbachia* infection following invasion into *Aedes aegypti* populations. PLoS Neglect Trop D 8: e3115.

Hoffmann, A.A., Montgomery, B.L., Popovici, J., Iturbe-Ormaetxe, I., Johnson, P.H., Muzzi, F. et al. (2011) Successful establishment of *Wolbachia* in *Aedes* populations to suppress dengue transmission. Nature 476: 454–457.

Huang, B.X., Yang, Q., Hoffmann, A.A., Ritchie, S.A., van den Hurk, A.F., and Warrilow, D. (2020) *Wolbachia* genome stability and mtDNA variants in *Aedes aegypti* field populations eight years after release. Iscience 23: 101572.

Ilinsky, Y. (2013) Coevolution of *Drosophila melanogaster* mtDNA and *Wolbachia* Genotypes. PLoS One 8: e54373.

Indriani, C., Tantowijoyo, W., Rances, E., Andari, B., Prabowo, E., Yusdi, D. et al. (2020) Reduced dengue incidence following deployments of *Wolbachia*-infected *Aedes aegypti* in Yogyakarta, Indonesia: a quasi-experimental trial using controlled interrupted time series analysis. Gates Open Res 4: 50.

Jasper, M.E., Yang, Q., Ross, P.A., Endersby-Harshman, N., Bell, N., and Hoffmann, A.A. (2019) A LAMP assay for the rapid and robust assessment of *Wolbachia* infection in *Aedes aegypti* under field and laboratory conditions. PLoS One 14: e0225321.

Jimenez, N.E., Gerdtzen, Z.P., Olivera-Nappa, A., Salgado, J.C., and Conca, C. (2019) A systems biology approach for studying *Wolbachia* metabolism reveals points of interaction with its host in the context of arboviral infection. PLoS Neglect Trop D 13: e0007678.

Johanowicz, D.L., and Hoy, M.A. (1998) Experimental induction and termination of non-reciprocal reproductive incompatibilities in a parahaploid mite. Entomol Exp Appl 87: 51–58.

Kong, J.D., Axford, J.K., Hoffmann, A.A., and Kearney, M.R. (2016) Novel applications of thermocyclers for phenotyping invertebrate thermal responses. Methods Ecol Evol 7: 1201–1208.

Kriesner, P., Conner, W.R., Weeks, A.R., Turelli, M., and Hoffmann, A.A. (2016) Persistence of a *Wolbachia* infection frequency cline in *Drosophila melanogaster* and the possible role of reproductive dormancy. Evolution 70: 979–997.

Kyle, J.L., and Harris, E. (2008) Global Spread and Persistence of Dengue. Annu Rev Microbiol 62: 71–92.

Lau, M.J., Ross, P.A., and Hoffmann, A.A. (2021) Infertility and fecundity loss of *Wolbachia*-infected *Aedes aegypti* hatched from quiescent eggs is expected to alter invasion dynamics. PLoS Neglect Trop D 15: e0009179.

Lau, M.J., Ross, P.A., Endersby-Harshman, N.M., and Hoffmann, A.A. (2020) Impacts of low temperatures on *Wolbachia* (Rickettsiales: Rickettsiaceae)-infected *Aedes aegypti* (Diptera: Culicidae). J Med Entomol 57: 1567–1574.

Lee, S.F., White, V.L., Weeks, A.R., Hoffmann, A.A., and Endersby, N.M. (2012) High-throughput PCR assays to monitor *Wolbachia* infection in the dengue mosquito *(Aedes aegypti)* and *Drosophila simulans*. Appl Environ Microb 78: 4740–4743.

Lewis, P.O. (2001) A likelihood approach to estimating phylogeny from discrete morphological character data. Syst Biol 50: 913–925.

Li, H. (2013) Aligning sequence reads, clone sequences and assembly contigs with BWA-MEM. arXiv:1303.3997.

Li, H., Handsaker, B., Wysoker, A., Fennell, T., Ruan, J., Homer, N. et al. (2009) The sequence alignment/map format and SAMtools. Bioinformatics 25: 2078–2079.

Lu, P., Bian, G.W., Pan, X.L., and Xi, Z.Y. (2012) *Wolbachia* induces density-dependent inhibition to dengue virus in mosquito cells. PLoS Neglect Trop D 6: e1754.

McMeniman, C.J., Lane, R.V., Cass, B.N., Fong, A.W.C., Sidhu, M., Wang, Y.F., and O’Neill, S.L. (2009) Stable introduction of a life-shortening *Wolbachia* infection into the mosquito *Aedes aegypti*. Science 323: 141–144.

Miller, M.A., Pfeiffer, W., and Schwartz, T. (2010) “Creating the CIPRES science gateway for inference of large phylogenetic trees” in proceedings of the gateway computing environments workshop (GCE), 14 Nov. New Orleans LA: 1–8.

Moreira, L.A., Iturbe-Ormaetxe, I., Jeffery, J.A., Lu, G.J., Pyke, A.T., Hedges, L.M. et al. (2009) A *Wolbachia* symbiont in *Aedes aegypti* limits infection with dengue, chikungunya, and plasmodium. Cell 139: 1268–1278.

Nazni, W.A., Hoffmann, A.A., NoorAfizah, A., Cheong, Y.L., Mancini, M.V., Golding, N. et al. (2019) Establishment of *Wolbachia* strain *w*AlbB in Malaysian populations of *Aedes aegypti* for dengue control. Curr Biol 29: 4241–4248.

Ng, L.C., and Project Wolbachia-Singapore Consortium. (2021) *Wolbachia*-mediated sterility suppresses *Aedes aegypti* populations in the urban tropics. medRxiv 2021.06.16: 21257922.

Olsen, K., Reynolds, K.T., and Hoffmann, A.A. (2001) A field cage test of the effects of the endosymbiont *Wolbachia* on *Drosophila melanogaster*. Heredity 86: 731–737.

Osborne, S.E., Iturbe-Ormaetxe, I., Brownlie, J.C., O’Neill, S.L., and Johnson, K.N. (2012) Antiviral protection and the importance of *Wolbachia* density and tissue tropism in *Drosophila simulans*. Appl Environ Microb 78: 6922–6929.

Pichon, S., Bouchon, D., Cordaux, R., Chen, L.M., Garrett, R.A., and Greve, P. (2009) Conservation of the type IV secretion system throughout *Wolbachia* evolution. Biochem Bioph Res Co 385: 557–562.

Pinto, S.B., Riback, T.I.S., Sylvestre, G., Costa, G., Peixoto, J., Dias, F.B.S. et al. (2021) Effectiveness of *Wolbachia*-infected mosquito deployments in reducing the incidence of dengue and other *Aedes-*borne diseases in Niteroi, Brazil: A quasi-experimental study. PLoS Neglect Trop D 15: e0009556.

Reed, L.J., and Muench, H. (1938) A simple method of estimating fifty per cent endpoints. American journal of epidemiology 27: 493–497.

Rice, D.W., Sheehan, K.B., and Newton, I.L.G. (2017) Large-scale identification of *Wolbachia* pipientis effectors. Genome Biol Evol 9: 1925–1937.

Richardson, M.F., Weinert, L.A., Welch, J.J., Linheiro, R.S., Magwire, M.M., Jiggins, F.M., and Bergman, C.M. (2012) Population genomics of the *Wolbachia* endosymbiont in *Drosophila melanogaster*. PLoS Genet 8:e1003129.

Riegler, M., Sidhu, M., Miller, W.J., and O’Neill, S.L. (2005) Evidence for a global *Wolbachia* replacement in *Drosophila melanogaster*. Curr Biol 15: 1428–1433.

Ritchie, S.A. (2018) *Wolbachia* and the near cessation of dengue outbreaks in Northern Australia despite continued dengue importations via travellers. J Travel Med 25: tay084.

Ross, P.A., and Hoffmann, A.A. (2018) Continued susceptibility of the *w*Mel *Wolbachia* infection in *Aedes aegypti* to heat stress following field deployment and selection. Insects 9: 78.

Ross, P.A., Endersby, N.M., and Hoffmann, A.A. (2016) Costs of three *Wolbachia* infections on the survival of *Aedes aegypti* larvae under starvation conditions. PLoS Neglect Trop D 10: e0004320.

Ross, P.A., Turelli, M., and Hoffmann, A.A. (2019a) Evolutionary ecology of *Wolbachia* releases for disease control. Annu Rev Genet 53: 93–116.

Ross, P.A., Ritchie, S.A., Axford, J.K., and Hoffmann, A.A. (2019b) Loss of cytoplasmic incompatibility in *Wolbachia*-infected *Aedes aegypti* under field conditions. PLoS Negl Trop Dis 13: e0007357.

Ross, P.A., Wiwatanaratanabutr, I., Axford, J.K., White, V.L., Endersby-Harshman, N.M., and Hoffmann, A.A. (2017) *Wolbachia* infections in *Aedes aegypti* differ markedly in their response to cyclical heat stress. PLoS Pathog 13: e1006006.

Ross, P.A., Gu, X.Y., Robinson, K.L., Yang, Q., Cottingham, E., Zhang, Y.F. et al. (2021) A *w*AlbB *Wolbachia* transinfection displays stable phenotypic effects across divergent *Aedes aegypti* mosquito backgrounds. Appl Environ Microb 87: e01264–01221.

Ryan, P.A., Turley, A.P., Wilson, G., Hurst, T.P., Retzki, K., Brown-Kenyon, J. et al. (2019) Establishment of *w*Mel *Wolbachia* in *Aedes aegypti* mosquitoes and reduction of local dengue transmission in Cairns and surrounding locations in northern Queensland, Australia. Gates Open Res 3: 1547.

Stamatakis, A. (2014) RAxML version 8: a tool for phylogenetic analysis and post-analysis of large phylogenies. Bioinformatics 30: 1312–1313.

Staunton, K.M., Yeeles, P., Townsend, M., Nowrouzi, S., Paton, C.J., Trewin, B. et al. (2019) Trap location and premises condition influences on *Aedes aegypti* (Diptera: Culicidae) catches using biogents sentinel traps during a ‘Rear and Release’ program: implications for designing surveillance programs. J Med Entomol 56: 1102–1111.

Stouthamer, R., Luck, R.F., and Hamilton, W.D. (1990) Antibiotics cause parthenogenetic Trichogramma (Hymenoptera, Trichogrammatidae) to revert to sex. P Natl Acad Sci USA 87: 2424–2427.

Suvakov, M., Panda, A., Diesh, C., Holmes, I., and Abyzov, A. (2021) CNVpytor: a tool for copy number variation detection and analysis from read depth and allele imbalance in whole-genome sequencing. Gigascience 10: giab074.

Tantowijoyo, W., Andari, B., Arguni, E., Budiwati, N., Nurhayati, I., Fitriana, I. et al. (2020) Stable establishment of *w*Mel *Wolbachia* in *Aedes aegypti* populations in Yogyakarta, Indonesia. PLoS Neglect Trop D 14: e0008157.

Trpis, M., Perrone, J.B., Reissig, M., and Parker, K.L. (1981) Control of cytoplasmic incompatibility in the *Aedes scutellaris* complex. J Hered 72: 313–317.

Truitt, A.M., Kapun, M., Kaur, R., and Miller, W.J. (2019) *Wolbachia* modifies thermal preference in *Drosophila melanogaster*. Environ Microbiol 21: 3259–3268.

Utarini, A., Indriani, C., Ahmad, R.A., Tantowijoyo, W., Arguni, E., Ansari, M.R. et al. (2021) Efficacy of *Wolbachia*-infected mosquito deployments for the control of dengue. New Engl J Med 384: 2177–2186.

van den Hurk, A.F., Hall-Mendelin, S., Pyke, A.T., Frentiu, F.D., McElroy, K., Day, A. et al. (2012) Impact of *Wolbachia* on infection with chikungunya and yellow fever viruses in the mosquito vector *Aedes aegypti*. PLoS Negl Trop Dis 6: e1892.

Veneti, Z., Clark, M.E., Zabalou, S., Karr, T.L., Savakis, C., and Bourtzis, K. (2003) Cytoplasmic incompatibility and sperm cyst infection in different *Drosophila-Wolbachia* associations. Genetics 164: 545–552.

Versace, E., Nolte, V., Pandey, R.V., Tobler, R., and Schlotterer, C. (2014) Experimental evolution reveals habitat-specific fitness dynamics among *Wolbachia* clades in *Drosophila melanogaster*. Mol Ecol 23: 802–814.

Walker, T., Johnson, P.H., Moreira, L.A., Iturbe-Ormaetxe, I., Frentiu, F.D., McMeniman, C.J. et al. (2011) The *w*Mel *Wolbachia* strain blocks dengue and invades caged *Aedes aegypti* populations. Nature 476: 450–453.

Wang, G.H., Gamez, S., Raban, R.R., Marshall, J.M., Alphey, L., Li, M. et al. (2021) Combating mosquito-borne diseases using genetic control technologies. Nat Commun 12: 1–12.

Wood, D.E., Lu, J., and Langmead, B. (2019) Improved metagenomic analysis with Kraken 2. Genome Biol 20: 1–13.

Wu, M., Sun, L.V., Vamathevan, J., Riegler, M., Deboy, R., Brownlie, J.C. et al. (2004) Phylogenomics of the reproductive parasite *Wolbachia* pipientis *w*Mel: a streamlined genome overrun by mobile genetic elements. PLoS Biol 2: 327–341.

Xi, Z.Y., Khoo, C.C.H., and Dobson, S.L. (2005) *Wolbachia* establishment and invasion in an *Aedes aegypti* laboratory population. Science 310: 326–328.

Yen, P.S., and Failloux, A.B. (2020) A Review: *Wolbachia*-based population replacement for mosquito control shares common points with genetically modified control approaches. Pathogens 9: 404.

Zheng, X.Y., Zhang, D.J., Li, Y.J., Yang, C., Wu, Y., Liang, X. et al. (2019) Incompatible and sterile insect techniques combined eliminate mosquitoes. Nature 572: 56–61.

